# Liquid-liquid phase separation of full-length prion protein initiates conformational conversion *in vitro*

**DOI:** 10.1101/2020.01.25.919340

**Authors:** Hiroya Tange, Daisuke Ishibashi, Takehiro Nakagaki, Yuzuru Taguchi, Yuji O. Kamatari, Hiroki Ozawa, Noriyuki Nishida

## Abstract

Prion diseases are characterized by accumulation of amyloid fibrils. The causative agent is an infectious amyloid that is comprised solely of misfolded prion protein (PrP^Sc^). Prions can convert PrP^C^ to proteinase-resistant PrP (PrP-res) *in vitro*; however, the intermediate steps involved in the spontaneous conversion remain unknown. We investigated whether recombinant prion protein (rPrP) can directly convert into PrP-res *via* liquid-liquid phase separation in the absence of PrP^Sc^. We found that rPrP underwent liquid-liquid phase separation at the interface of the aqueous two-phase system (ATPS) of polyethylene glycol (PEG) and dextran, whereas single-phase conditions were not inducible. Fluorescence recovery assay after photobleaching revealed that the liquid-solid phase transition occurred within a short time. The aged rPrP-gel acquired proteinase-resistant amyloid accompanied by β-sheet conversion, as confirmed by western blotting, Fourier transform infrared spectroscopy, and Congo red staining. The reactions required both the N-terminal region of rPrP (amino acids 23-89) and kosmotropic salts, suggesting that the kosmotropic anions may interact with the N-terminal region of rPrP to promote liquid-liquid phase separation. Thus, structural conversion *via* liquid–liquid phase separation and liquid–solid phase transition are intermediate steps in the conversion of prions.

## Introduction

Transmissible spongiform encephalopathies (TSEs), also called prion diseases, are infectious and fatal neurodegenerative diseases with rapidly progressive dementia, such as Creutzfeldt–Jakob disease (CJD) in humans (1). TSEs are characterized by the accumulation of misfolded prion protein (PrP^Sc^), which is spontaneously converted from normal prion protein (PrP^C^). PrP^C^ is well preserved among mammalian species and is particularly expressed in the neurons and tethered to the cell membrane *via* the glycosylphosphatidylinositol (GPI) anchor (2). The protein-only hypothesis proposes that the infectious agent, prion, is solely composed of PrP^Sc^. The main biochemical characteristics of PrP^Sc^ are that it is a protease K-resistance fragment (PrP-res) and it has seeding activity to convert PrP^C^ into itself (PrP^Sc^) (3-5). This conversion process presumably proceeds *via* direct interaction between PrP^C^ and PrP^Sc^ (6). Several studies have attempted to generate artificial PrP^Sc^, and the amplification of PrP^Sc^ *in vitro* has been successfully demonstrated using intermittent ultrasonication on the brain homogenates, called protein misfolding cyclic amplification (PMCA) (7,8). Not only sonication but also shaking of the protein solution can promote *in vitro* amyloid formation. The quaking-induced conversion (QuIC) assay is now widely used to detect trace amounts of PrP^Sc^ in cerebrospinal fluid using rPrP as a substrate (9). These lines of experimental evidence suggest that rPrP can be converted to proteinase K-resistant amyloid (rPrP-res) in the presence of PrP^Sc^ with the provision of kinetic energy. However, to explain spontaneous generation and to generate artificial prions, the spontaneous misfolding process from rPrP to rPrP-res in the absence of PrP^Sc^ needs to be elucidated.

Recently, proteins with intrinsically disordered regions (IDRs) have been shown to undergo liquid phase separation in the cytoplasm and form membrane-less organelles such as stress granules (10). In the liquid phase, IDRs assemble to form a cross-β sheet structure. This phenomenon has been associated with the development of neurodegenerative diseases, including Tau protein in Alzheimer’s disease and FUS in amyotrophic lateral sclerosis, which are caused by pathogenic amyloids (11,12). Therefore, the aberrant phase transition of amyloidogenic proteins may facilitate pathological amyloid synthesis.

The N-terminal of PrP^C^ is an IDR comprised of 5 repeats of proline/ glycine−rich sequences, which are called octapeptide repeats. Therefore, to elucidate the spontaneous process involved in the conversion of PrP^C^ into PrP^Sc^, we examined whether rPrP can convert into rPrP−res or PrP^Sc^ *via* liquid-liquid phase separation, without the use of kinetic energy. In this study, we found that liquid-liquid phase separation and liquid-solid phase transition of rPrP require the interaction between the N-terminal region and kosmotropic anions. Furthermore, the rPrP in gels acquired the features of PrP-res with β-sheet-rich structure and protease-K resistance. These results suggest that the liquid-liquid phase separation and liquid-solid phase transition can initiate spontaneous conformational conversion of rPrP to PrP-res without the use of kinetic energy, and that the interaction between kosmotropic anions and N-terminal region of PrP plays a key role in the conformational conversion process in prion diseases.

## Materials and Methods

### Protein expression and purification

We prepared 3 rPrPs: full-length human PrP (residues: 23–231), truncated human PrP (residues: 90–231), and Mo-rPrP (residues: 23–231). All constructs were expressed in *Escherichia coli* strain DH5α. The expression and purification of rPrPs were performed as previously described (9,47). After purification, each protein solution was frozen at −80°C in 150 µL aliquots, which were thawed for single use. Before using for any experiment, each protein solution was centrifuged at 15,000 rpm for 10 min at room temperature (28°C). To prepare labeled rPrP, Alexa Fluor 488 Microscale Protein Labeling Kit (A30006, Invitrogen, Carlsbad, CA, USA) was used. The procedure was performed in accordance with the manufacturer’s instructions.

### Disorder propensity and charge prediction

Disorder propensity was calculated using PrDOS (18), charge prediction was performed using EMBOSS (19,20), and the hydrophilic region was calculated using ProtScale (21). The amino acid sequence from Uniprot (P04156) was used.

### Droplet formation assay (polymer and salt solution preparation)

Polymer solutions were prepared from polyethylene glycol (PEG) (MW: 6000) (Wako, Osaka, Japan) and dextran (MW: 180,000) (Nacalai Chemical, Kyoto, Japan). Each component was dissolved in dH_2_O and prepared as 50% PEG and 25% dextran (w/v) and stored at 4°C in 1 mL aliquots. The phase diagram was created by direct observation of polymer droplets using differential interference contrast microscopy using (DIC). PEG-dextran polymer solutions were prepared as 1-10 % (wt/vol) of each polymer in 20 µL of solution. The polymer solutions were vigorously vortexed, and 5 µL of the solution was loaded onto a slide glass. For confocal microscopy observation with fluorescence, 0.01% rhodamine-PEG (#PG1-RB-5k, Nanocs, New York, NY, USA) was used. For salt solutions, we prepared 2 M stocks of NaCl, Na_2_S_2_O_3_, Na_2_CO_3_ (Wako, Osaka, Japan) Na_3_Citrate, Na_2_SO_4_, and (NH_4_)_2_SO_4_ (Nacalai Chemical, Kyoto, Japan). Each solution was stored at room temperature. To prepare the aqueous two−phase system (ATPS) solution, each polymer solution was mixed at concentrations ranging from 1.5% to 13.5% with 200 mM salt (final: 1–9 wt% of each polymer, PEG/ dextran with 120 mM salt). Then, the solution was pipetted well and vigorously vortexed. Experiments were performed on a scale of 50 µL (52.6 µl with Thioflavin T [ThT]); 2.6 µL of 1 mM ThT (final concentration: 50 µM) was added 30 µl of ATPS solution and then Next, 20 µl of rPrP solution (final concentration: 2–10 μM) was added to the ATPS solution and gently pipetted 10–15 times. The entire solution was applied to a glass slide or 96-well plate (#165305, Thermo Fisher Scientific, Waltham, MA, USA) for microscopic observation. Droplet observation was performed using confocal microscopy (#LMS700; Carl Zeiss, Oberkochen, Germany) and DIC microscopy (Axioskop2; Carl Zeiss, Oberkochen, Germany) with 20x and 40x objective lenses. To evaluate ThT fluorescence, Colibri 7 (Carl Zeiss, Oberkochen, Germany) was used as a luminous source at a wavelength of 485 nm. Images were acquired with exposures of 250 ms (low exposure), 500 ms, and 2000 ms (high exposure). The pH was adjusted using NaOH (1N) or HCl (1N) and confirmed by test paper. For droplet aging, the samples were applied to a 96-well plate or Eppendorf tube incubated at 37°C for 30 min to 72 h. All experiments were performed in triplicate.

### Congo red staining

The samples were incubated for 72 h at 37°C in the ATPS solution. After incubation, 200 µl of dH_2_O was added and pipetted well. The aged gels were collected by centrifugation at 15,000 rpm for 10 min at room temperature and were stained with 50 µL of 1% Congo red (#C8,445-3: Aldrich) solution for 30 min in an Eppendorf tube atroom temperature. After staining, the sample was centrifuged again under similar conditions, and the supernatant was discarded. The pellet was washed with 50 µL of dH_2_O by pipetting, centrifuged again under similar conditions, and the supernatant was discarded. The pellet was suspended in 20 µL of dH_2_O, and 5 µL was applied on a glass slide, followed by sealing with a cover glass. Microscopic observation was performed using a confocal microscope (Nikon, Tokyo, Japan) with a polarization filter. Images were acquired using the NIS-Elements C software.

### Fluorescence after photobleaching assay

FRAP was performed using the LMS700. Alexa488-labeled human rPrP was diluted 1:18 with native human rPrP (final: 13 µM). Bleaching was performed with 100% transmission of a 405, 488, or 500 nm laser. Pre-bleaching images were taken for 3s (1s frame rate, 3 frames), whereas post-bleaching images were acquired for the following 120 s (1s frame rate, 120 frames) and analyzed with ZEN. The samples named “0 min” were taken in less than 5 min, including the set up. The sizes of the bleached area and background area were set in the first experiment. For each image, the bleached region and background region were calculated using ZEN, and the background was subtracted during analysis.

### Sarkosyl and proteinase K treatments

Sarkosyl and PK were purchased from Sigma-Aldrich. For sarkosyl treatment, the sample was incubated in the ATPS solution at 37°C for 30 min, and 200 µL of dH_2_O was added to the sample and pipetted well. The entire solution was centrifuged at 15,000 rpm for 10 min at room temperature. Supernatant-1 (S1) and Pellet-1 (P1) were collected. P1 was suspended in 25 µL of dH_2_O or 1% sarkosyl and then incubated at 37°C for 10 min. After incubation, samples were centrifuged at 15,000 rpm for 30 min at room temperature and then Supernatant-2 (S2) and Pellet-2 (P2) were retrieved. The PK solution (10 µg/ml) was prepared in dH_2_O. The samples were incubated at 37°C for 72 h in an Eppendorf tube. As a negative control, the solution containing the same amount of rPrP was treated with the PK solution. The samples and PK solution were mixed by pipetting, applied to a 96-well plate, and incubated at 37°C. DIC microscopy was performed at the beginning of the reaction (0 min) and the end of incubation (30 min). Samples were retrieved from the 96-well plate, and each well was washed with 100 µL of dH_2_O. The entire sample was collected in an Eppendorf tube and centrifuged at 15,000 rpm for 10 min at room temperature. The supernatant and pellet were collected. In both experiments, the supernatant was denatured with 6x SDS sample buffer (50 mM Tris-HCl pH 6.8, 5% glycerol, 1.6% SDS, and 100 mM dithiothreitol). The pellet was then resuspended in 1x SDS buffer and boiled at 95°C for 10 min for SDS-PAGE.

### Immunoblotting

Samples were loaded to 18% acrylamide gel for SDS-PAGE and then transferred to Poly Vinylidene Di-Fluoride membrane. The membrane was blocked using 5%(w/v) skim milk with TBST (10 mM Tris-HCl pH 7.8, 100 mM NaCl, 0.1% Tween 20) at RT for 1 h. To detect PrP, the membrane was incubated with primary antibody R20 (1:1000 diluted with 1% skim milk) for 1 h at RT (48). Horseradish peroxidase-conjugated anti-rabbit IgG (1:10,000, GE Healthcare Life Sciences, Fairfield, CT, USA) was used as the secondary antibody. Protein bands were visualized using Clarity Western ECL substrate (Bio-Rad, Hercules, CA, USA). Band intensity was quantified using ImageJ.

### Quantifying ThT fluorescence

Fluorescence intensity was quantified with FLUOstar Omega (BMG Labtech, Ortenberg, Germany) in a 96-well plate with a spiral scan. The 96-well plate was covered with sealing tape (#J676060, Greiner, Kremsmünster, Austria), incubated at 37°C, and monitored by the bottom reading of the fluorescence intensity every hour up to 48 h using monochromators or filters with wavelengths of 448 nm (excitation) and 482 nm (emission).

### Fourier transform infrared spectroscopy analysis

Fourier transform infrared spectroscopy (FTIR) spectra were measured using a JASCO FT/IR-4700ST with attenuated total reflection. Five microliters of the sample was loaded onto the grid. To prepare the sample for FTIR, we first prepared a 30x concentrated sample (aged for 72 h) from 1.5 mL scale to 50 µL. Aged gel was collected by centrifugation, as described above, and suspended in dH_2_O.

## Results

### rPrP undergoes liquid–liquid phase separation in ATPS

In general, polymers such as PEG or dextran are used to induce liquid-liquid phase separation of proteins as crowding agents (13). First, we tried with a single polymer solution; however, rPrP did not undergo liquid-liquid phase separation but resulted in salting out with both PEG and dextran at concentrations greater than 10% (Sup Fig. 1A). We applied ATPS, which is composed of PEG and dextran, because the interface of ATPS may function like a cellular surface (14). The droplets appeared at the interface of the polymer fractions immediately after mixing 10 µM rPrP with an ATPS mixture containing sodium thiosulfate (Na_2_S_2_O_3_). We tested combinations of various concentrations of the polymers and investigated where ATPS could form an interface (15) (Fig. 1A, B). Below the binodal curve, no droplet was formed at the interface of the ATPS. Under such conditions, rPrP precipitated as amorphous aggregates at the bottom of wells after 24 h of incubation (Sup Fig. 1B). With 6%/ 6% PEG/ dextran, the spherical droplets were observed at the interface of ATPS and bottom of the well, and the amorphous aggregates were visualized by ThT. The spherical droplets appeared even more efficiently with 9%/ 9% PEG/ dextran (Fig. 1B, Sup Fig. 1B). Quantification of ThT fluorescence intensity showed that 9%/ 9% of PEG/ dextran had the highest fluorescence intensity after 24 h of incubation (Supplementary Fig. 1C). Therefore, we set the experimental conditions of 9%/ 9% PEG/dextran with 120 mM Na_2_S_2_O_3_ in the following experiments, unless otherwise noted. ThT-positive aggregates appeared to correlate with the concentration of rPrP for up to 6 µM; spherical droplets with clear ThT fluorescence appeared from 8 µM rPrP and were most efficient at 10 µM of rPrP (Supplementary Fig. 2A). The fluorescence intensity was significantly high in the presence of 10 µM of rPrP (Sup Fig. 2B). To confirm if the droplets consisted of rPrP, we performed a similar experiment with Alexa 488−labeled rPrP and found that the fluorescence was equally distributed in all the droplets (Fig. 1C). The droplets could be visualized by ThT immediately after their formation, suggesting that β-sheet formation of rPrP was initiated inside the fresh formed droplets.

**Fig. 1.**
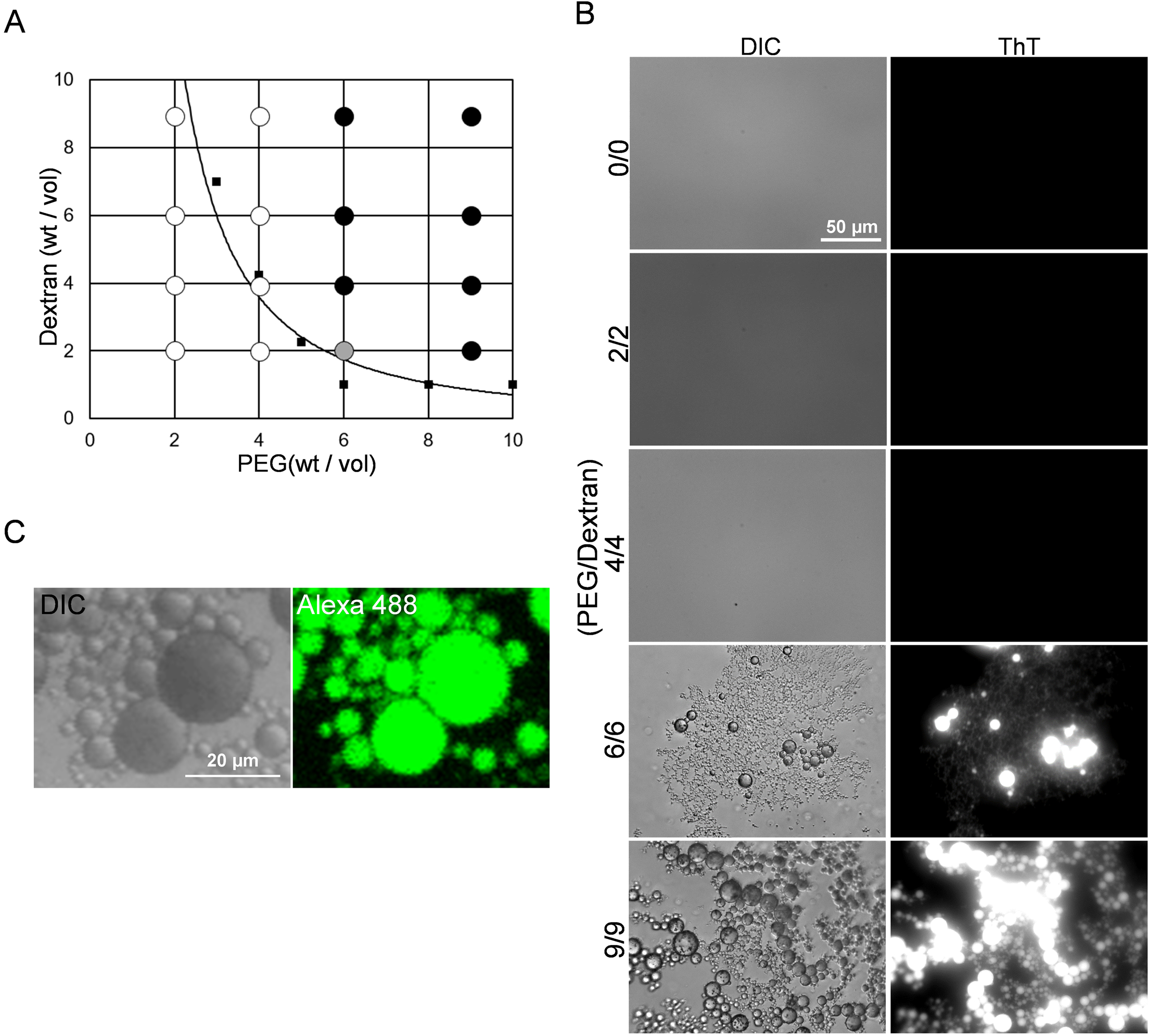
rPrP undergoes liquid-liquid phase separation in ATPS. (A) Phase diagram of an ATPS (PEG/ dextran). The binodal curve (solid line) is drawn with approximation (R^2^=0.8626). Black dots: rPrP fully undergo liquid phase separation. Gray dots: rPrP underwent LLPS with aggregation. White dots: rPrP fully aggregated. Square dots: Average of clouding point. N=3 (B) Differential interference contrast (DIC) and fluorescence microscopic images of droplets in the interface of PEG/ dextran after 24 h of incubation. Bar: 50 µm. (C) Confocal microscopic images of rPrP droplets with Alexa 488 labeled rPrP (1:18). Bar: 20 µm.

### Kosmotropic anion species induce droplet formation

We next investigated the influence of the salt types on droplet formation, and screened various salts according to the Hofmeister series. Sodium salts, such as NaCl, Na_2_S_2_O_3_, Na_2_CO_3_, Na_3_Citrate, Na_2_SO_4_, and (NH_4_)_2_SO_4_. Na_2_SO_4_, Na_3_Citrate, and (NH_4_)_2_SO_4_ were tested. We found that the kosmotropic salts induced droplet formation, whereas NaCl and Na_2_CO_3_ did not (Sup Fig. 3A). The droplets in the Na_2_CO_3_ samples had almost no fluorescence intensity because the alkaline conditions caused by Na_2_CO_3_ affected the ThT, resulting in loss of fluorescence ability (16). These results suggest that kosmotropic anions have a strong influence on the rPrP droplet formation. We next examined the influence of pH under conditions of 9%/ 9% of PEG/ dextran with 120 mM Na_2_S_2_O_3_. At pH 4, a small number of spherical droplets were observed, but most of them formed ThT-positive, granule–like aggregates with a low value of circularity,. These granule-like aggregates did not fuse with each other. Among the conditions we tested, the droplets were most efficiently formed at neural pH, although we could not fully evaluate the formation efficiencies at pH 12 due to loss of ThT fluorescence (Sup Fig. 3C). We confirmed that 120 mM Na_2_S_2_O_3_ and neural pH were the optimal conditions for our experiments.

### The droplets of rPrP undergo liquid-solid phase transition

To investigate the properties of the droplets, we continuously observed their behavior. The fresh droplets floating at the interface seamlessly fused with each other, suggesting that the droplets were in the liquid phase (Fig. 2A). Further, rPrP immediately condensed to form droplets at the interface of PEG/ dextran by adding Na_2_S_2_O_3_. In addition, fluorescence-labeled PEG colocalized with rPrP in the droplets, suggesting that PEG was bound to rPrP (Fig. 2B). Next, we analyzed the fluorescence recovery after photobleaching of the droplets, before and after 1 h of incubation. Fresh droplets, immediately after liquid-liquid phase separation (0 min), showed full recovery of the intensity within 60 s after photobleaching, whereas the droplets incubated for 1 h at 37°C showed no recovery throughout the observation period (Fig. 2C, D), suggesting that the droplets of rPrP underwent liquid–solid phase transition and became rPrP-gels.

**Fig. 2.**
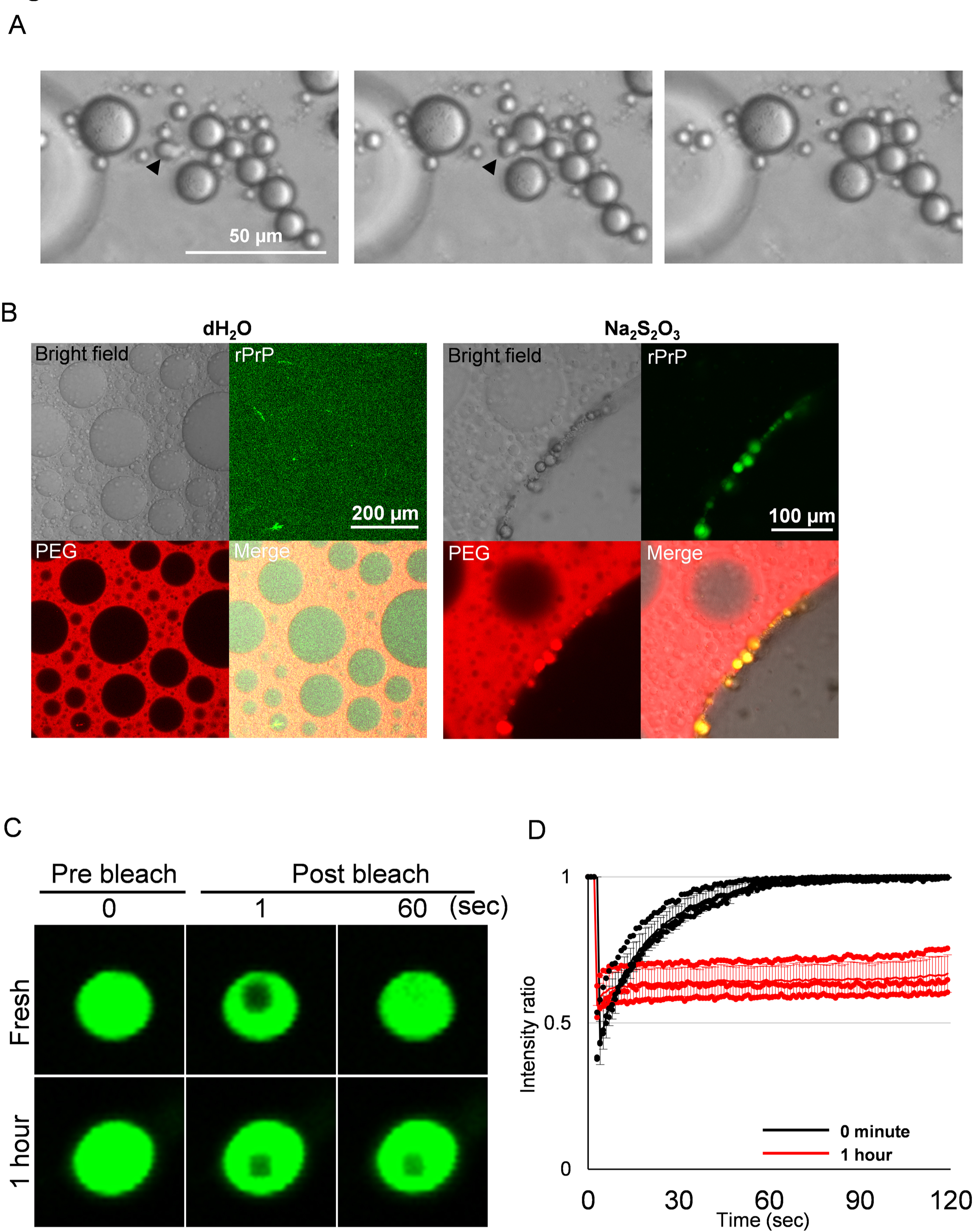
rPrP droplets undergo liquid-solid phase transition. (A) Consecutive images of droplets seamlessly combining with each other. Black arrowhead indicates a droplet uniting with another one. (B) Confocal microscopic images of distribution of PEG (0.01% of Rhodamine-PEG: red) and rPrP (1:18 of Alexa488-labeled rPrP: green). Left panel: ATPS/ rPrP solution with dH_2_O. Bar: 200 µM. Right panel: ATPS/ rPrP solution with 120 mM of sodium thiosulfate. Bar: 100 µm. Both images were acquired immediately after mixing the polymer and rPrP solution. (C) Confocal microscopic images from FRAP experiment of a fresh (0 min) droplet (top) and a droplet incubated for 1 h (bottom). Bar: 5 µm. (D) Fluorescence recovery after photobleaching (FRAP) curves from flesh droplets (0 min) and droplets incubated for 1 h regarding in 2C. Each dot indicates a value measured in 3 independent experiments. Line indicates average value. Bar: SD. N=3.

### The N-terminal region of rPrP (residues 23–89) drives liquid-liquid phase separation and liquid-solid phase transition

The N-terminal region of PrP^C^ is known to be an IDR, whereas its C-terminal consists of stable secondary structures with 3 α-helices, as revealed by Nuclear Magnetic Resonance study, consistent with the prediction result from PrDOS (17,18). Under biological conditions, PrP^C^ is not phosphorylated or methylated but is a GPI-anchored protein with two glycosylation sites (Fig. 3A). To determine whether the IDRs of rPrP influence liquid-liquid phase separation, we first calculated its disordered propensity, hydrophobicity, and electric charge (Fig. 3B). This region coincides with the positively charged region predicted by EMBOSS and the hydrophilic region calculated from Protscale (19-21). To elucidate the role of the N-terminal region, we compared the behavior of the full-length rPrP and N-terminally truncated mutant, rPrP Δ (23–89) in ATPS. We found that rPrP Δ (23–89) did not increase the fluorescence intensity even with Na_2_S_2_O_3_, but formed slightly ThT-positive aggregates at the interface. These aggregates showed no increase in ThT fluorescence throughout the observation period of up to 48 h, whereas the droplet of full-length rPrP increased the fluorescence intensity over time (Fig. 3C-E). Furthermore, the fluorescence intensity was significantly higher than that of rPrP Δ (23–89) with Na_2_S_2_O_3_ at 1 h and became more striking after 48 h (Fig. 3F, G). Full-length mouse rPrP (Mo−rPrP-residues: 23–231) also showed similar results (Supplementary Fig. 4 A, B).

**Fig. 3.**
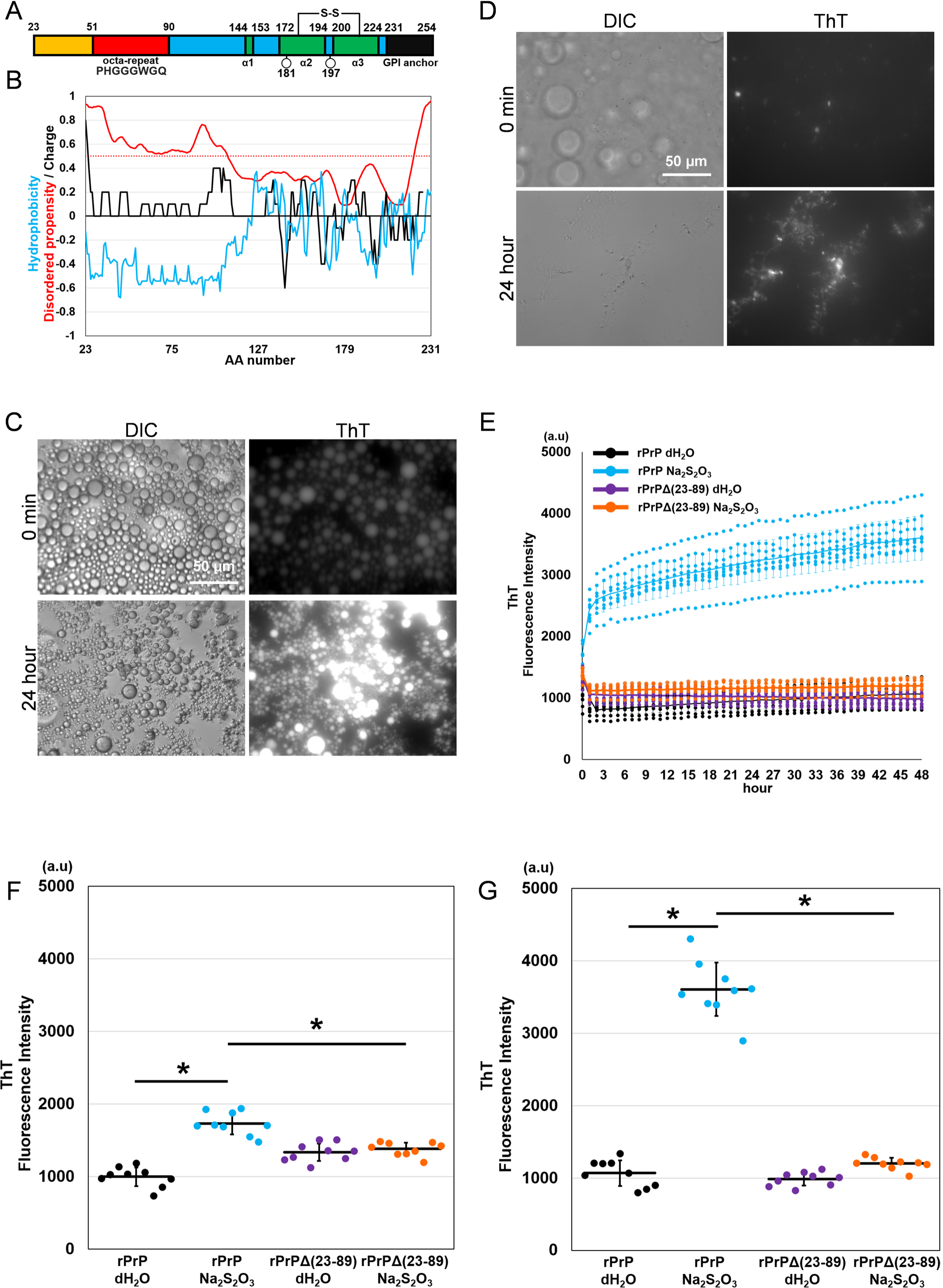
The N-terminal of PrP is essential for droplet formation and maturation. (A) Schematic diagram of human rPrP residues 23-231. Red indicates octapeptide repeats region. Green indicates alpha helix regions. Circles indicates glycosylation sites. (B) Calculation of hydrophobicity, disordered propensity, and electric charge of human prion protein using Protscale, PrDOS, and EMBOSS. Blue line: hydrophobicity calculated by Protscale. Black line: electric charge calculated by EMBOSS. Red line: disordered propensity calculated by PrDOS. Red dot line: threshold of disordered propensity (FP: 5%). (C) DIC and fluorescence microscopic images of rPrP: Sodium thiosulfate at 0 min and 24 h. Even very small droplets (< 5 µm) with no apparent ThT fluorescence at 0 min could be clearly identified after 24 h of incubation. (D) DIC and fluorescence microscopic images of rPrP Δ (23-89): Na_2_S_2_O_3_ at 0 min and 24 h, acquired h long exposure. (E) ThT fluorescence intensity of rPrP and rPrP Δ (23-89) measured in 48 h. Each dot represents a value measured in 3 independent experiments. Line indicates average of each group. (F, G) Quantification of ThT fluorescence intensity at 0 min and 48. *P<0.0001. Bar represents SD. N=9. Statistical analysis was performed with one-way ANOVA, followed by the Tukey-Kramer test.

### Liquid-solid phase transition involves conformational conversion of the prion protein

Because the rapid growth of ThT fluorescence coincides with the timing of a liquid–solid phase transition (Fig. 2C, D), we postulated that the droplets rapidly became insoluble and eventually matured to acquire the properties of PrP^Sc^. To verify this, we incubated the droplets for 30 min and then collected them by centrifugation. The rPrP-gels were ThT-positive and did not dissolve in water (Fig. 4A). Subsequently, we resuspended the gels in 1% sarkosyl and reprecipitated them by centrifugation. Western blot analysis showed that rPrP was insoluble in 1% sarkosyl solution (Fig. 4B). There was no significant difference in the insoluble fraction (P2), with or without treatment (Fig. 4C). Next, we examined whether these PrPs in the gels acquired PK resistance. We aged rPrP-gels for 72 h and then digested them with PK. The appearance of aged gels remained unchanged after PK digestion (Fig. 4D). SDS−PAGE and western blotting of the aged gels collected by centrifugation showed that the aged droplets contained oligomers of rPrP, and 40% of rPrP remained undigested (Fig. 4E, F). A small PrP-res fragment was detected around 10-15 kDa, resembling the PMCA product (25). We could not disrupt the aged gel by sonication to improve the penetration of proteinase. We further attempted to confirm that the rPrP-gel was composed of amyloid. The aged gels stained positively with Congo red, although they did not show apple-green birefringence under cross-polarized light (Fig. 4G). A similar observation was reported in human amyloid spherulites composed of islet amyloid polypeptide in the pancreatic tissue of type 2 diabetes mellitus (22). However, the aged gel did not show the Maltese cross under cross-polarized light, which is a characteristic of spherulites. To analyze the secondary structure of the aged gel, we performed FTIR analysis. FTIR analysis showed that the aged gels had a distinctive peak at 1620 cm^−1^ in the β-sheet region of the second-derivative spectra, which shifted from 1651 cm^−1^ in the α−helix region from the native form of rPrP (Fig. 4H). Moreover, they were stable for months in water (data not shown). These results suggest that rPrP is converted into PrP-res inside the droplets, acquiring β-sheet structure, detergent-insolubility, and PK resistance.

**Fig. 4.**
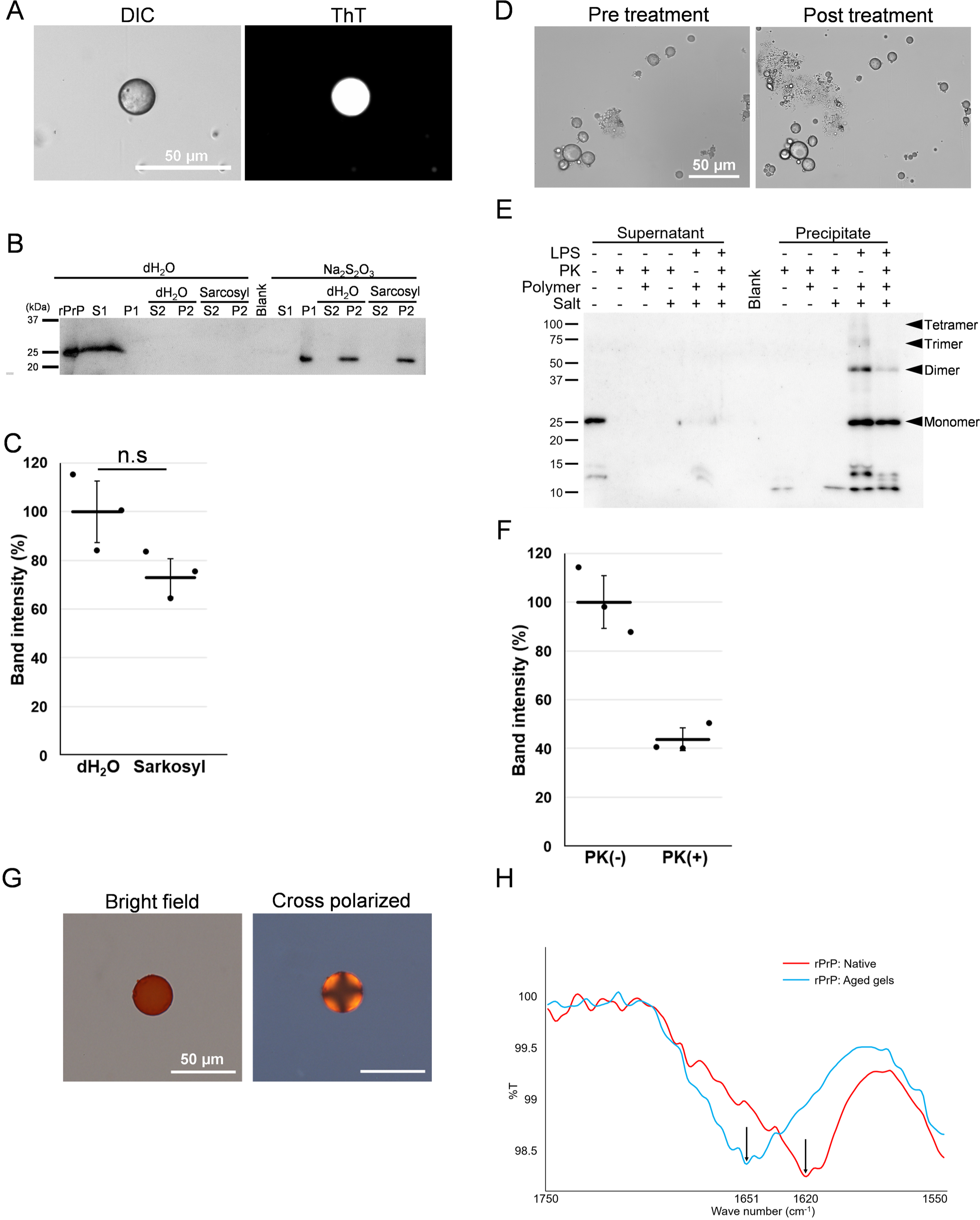
Biochemical analysis of the rPrP-gel. (A) DIC and fluorescence microscopic images of the rPrP-gel incubated for 30 min in ATPS, collected by centrifuge, and then applied into dH2O with 50 µm of ThT. Bar: 50 µm. (B) Western blotting of rPrP, with or without sarkosyl treatment. S1 and P1 were originally collected by centrifugation from the sample diluted with dH2O. S2 and P2 were collected from the P1 fraction treated with dH2O or sarkosyl. (C) Quantification of band intensity from P2 fraction of dH2O comparing with sarcosyl treatment (refer to Fig 4B).(D) DIC microscopic images of aged gels before (0 min) and after (30 min) PK treatment. (E) Quantification of band intensity of aged gels, with or without PK treatment (refer to Fig. 4D). (F) Western blotting of rPrP aged gels. “LPS” indicates that experiments were done under the condition of 9%/ 9% of PEG/ dextran, 120 mM of sodium thiosulfate and 10 µM of rPrP. “dH2O” indicates that experiments were done under the condition of 9%/ 9% of PEG/ dextran, 10 µM of rPrP without salt. “Polymer” and “Salt” indicates that the samples contain 9%/ 9% of PEG/ dextran, and 120 mM of sodium thiosulfate. “PK” indicates that samples were treated with 7.5 ug/ml of PK. Each dot represents a value measured from individual sample. Solid line indicates average. Bar: SD. N=3. Statistical analysis was performed with unpaired t-tests. “n.s.” means no significant difference. (G) Confocal microscopic images of aged rPrP-gels stained with Congo-red in bright field and cross polarized. Bar: 50 µm. (H) FTIR spectroscopic analysis of aged rPrP-gels and native rPrP. Blue line: native rPrP. Red line: Aged rPrP-gels. Arrows indicate the peak of each sample.

## Discussion

We have demonstrated that rPrP undergoes liquid-liquid phase separation at the ATPS interface. IDR in the N-terminal region of PrP^C^ (residues: 23-89) and kosmotropic anions in the ATPS were essential for the overall reaction. The rPrP liquid droplets subsequently showed liquid–solid phase transition within an hour, and the aged rPrP gels contained β-sheet-rich, sarkosyl-insoluble, and PK-resistant amyloid. These data suggest that PrP^C^ in the liquid phase can initiate spontaneous conversion to amyloids without the provision of kinetic energy or seeding.

ATPS has been used for a wide range of purposes, such as purification of enzymes, nucleic acids, and viruses for providing a gentle environment for biomolecules and stabilizing their structure (15,23). The partitioning behavior in ATPS has been well documented. In protein purification, monomeric IgG could be collected separately in the PEG-rich fraction because of its positive charge (24). Such convention by ATPS can facilitate interactions of the sequestered molecules, as demonstrated by DNA and actin fibers separately interacting and polymerizing inside the dextran phase, called cell-sized aqueous microdroplets, imitating cell-like crowded microenvironments (25). Similarly, our present results could be interpreted from a similar viewpoint. It is conceivable that rPrP, which has positive charges like IgG in the IDR, was sequestered to the PEG phase at first and then when it was sufficiently condensed, it formed a liquid phase on its own owing to the interactions between the IDRs. Similar to many other proteins known to undergo liquid-liquid phase separation, the region consists of 5 repeats of a glycine-rich motif and contains proline and aromatic amino acids, that is, tryptophan.

The kosmotropic anions have been shown to stabilize the structure of proteins to enhance amyloid formation *in vitro* (26,27). It has been shown that the efficiency of amyloid formation from prion protein is in accordance with the Hofmeister series (28,29). In addition, kosmotropic anions have been shown to drastically improve the detection limit of pathological amyloids, including prions (30). Therefore, kosmotropic anions may play a role in promoting structural stabilization of the proteins, facilitating their transformation to amyloids after the formation of droplets. However, further investigation is required to elucidate the role of kosmotropic anions from the viewpoint of the electrical effect.

It is unlikely that the fluorescence-labeled PEG that colocalized with the droplet of rPrP contribute directly to the reactions of rPrP in the liquid phase separation because it hardly affects the conversion properties of rPrP to PrP-res (31). Analogous to other proteins that undergo liquid-liquid phase separation, the proline and glycine-rich N-terminal IDR of rPrP are very flexible and multivalent because of the periodically located tryptophan residues; these features enable efficient intermolecular interactions and consequently liquid-liquid phase separation. The liquid-phase formation *via* IDR may subsequently facilitate interactions between the C-terminal regions, and finally evoke parallel β-sheet conversion of the entire molecule. Therefore, it does not require agitation to catalyze the reactions. This may be in contrast to the facilitation of conversion by mechanical agitation, where natively folded protein molecules at the air/ water interface are denatured and the forcefully exposed hydrophobic residues presumably enable efficient intermolecular binding, and eventually conversion (32-34).

Our view that the IDR of rPrP drives liquid-liquid phase separation and that liquid-solid phase transition is accompanied by the conversion of the C-terminal region is valid. It has been shown that IDR assembles protein molecules and forms a cross-β structure comprising stacks of short β-strands in the process of liquid-solid phase transition (35).

Similar to other proteins that are reported to undergo liquid-liquid phase separation, IDRs of PrP, that is, octapeptide (PHGGGWGQ) repeats, are very flexible and multivalent; thus, the octapeptide region quickly forms a short cross-β structure, as suggested by the ThT positivity at the very beginning of droplet formation. Inside the droplet, the flexible intermolecular interactions of rPrP through the octapeptide repeats maintain the C-terminal regions of rPrP in the proximity of each other until they are fully converted into β-sheet-rich structures. In addition, repeats of the motif have advantages in the liquid phase because of the high plasticity of intermolecular bindings under shear stress. In summary, we propose that (i) the N-terminal region with positive charges induces condensation of rPrP in the PEG phase, (ii) the charges are neutralized by the kosmotropic anion, inducing direct interaction of dipole (G,Q) and π-π stacking (W) of the octa-peptide region to form a short cross-β sheet structure (36), (iii) the molecular distance of the C-terminal region is reduced enabling them to become close to each other, leading to the polymerization and β-sheet conversion of the entire rPrP to the amyloid. (Fig 5A, B). This process may be similar to the *in silico* simulation model suggesting that the conversion process started from the N-terminal region (37).

**Fig. 5.**
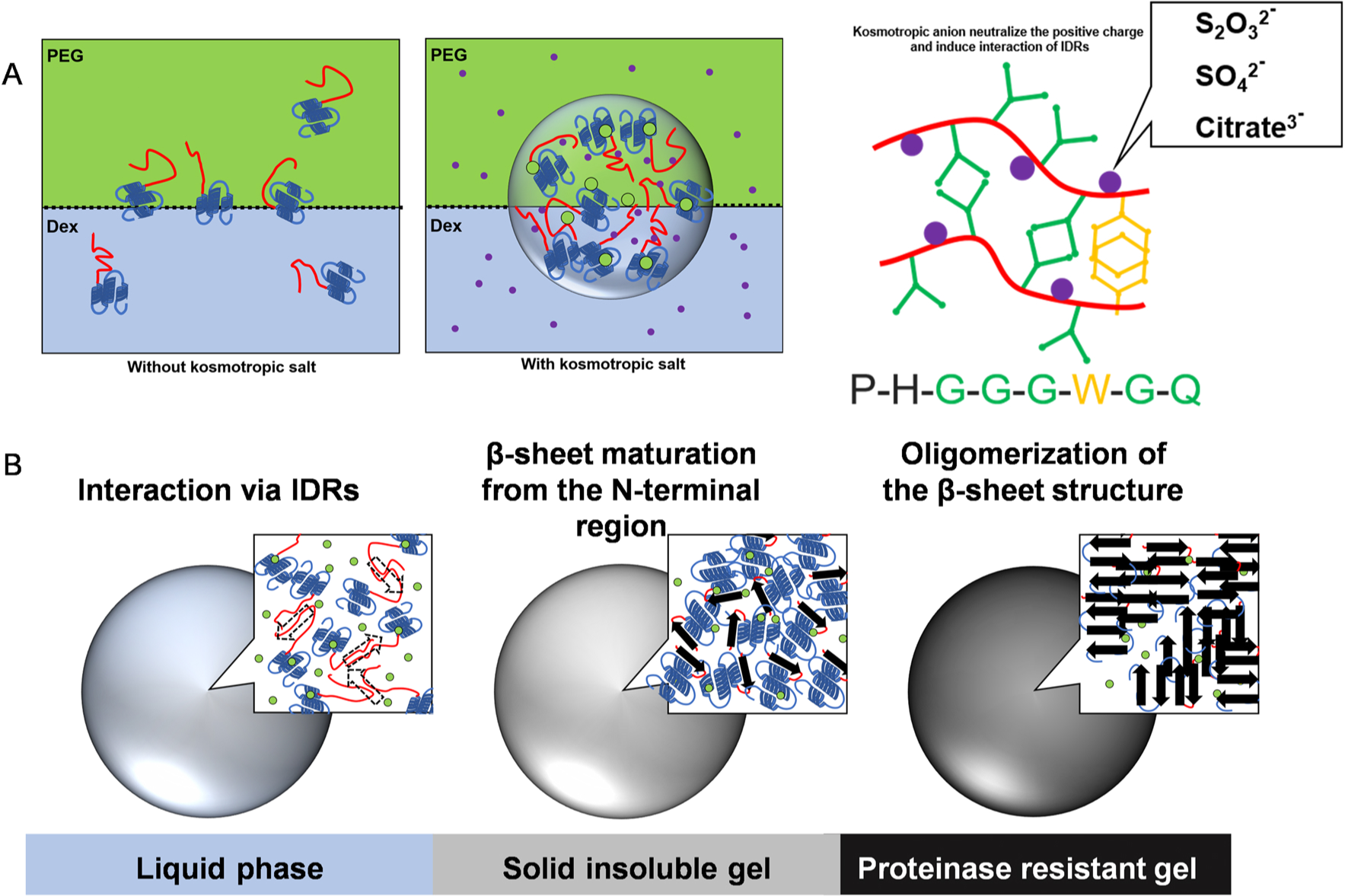
Hypothetical model for droplet formation and phase transition. (A) Left: Prion protein molecules are equally dissolved in PEG (yellow green) and dextran (blue) fractions without kosmotropic anions. Middle: Prion protein molecules assemble each other *via* IDRs and form droplets at the interface of ATPS by adding kosmotropic anions (purple dots). PEG may bind to prion protein but with no change in its structure. Right: Possible IDRs (red line) interaction inside droplets; Kosmotropic anions neutralize the positive charge of IDRs and induce interaction between them. Dipole–dipole interaction (green) of glycine (G) and glutamine (Q), and π–π interaction (yellow) of tryptophan (W) are expected to underlie in phase separation and transition (refer to (32]). (B) Hypothetical model of liquid-solid phase transition. In a fresh droplet, the IDRs of prion protein may construct a cross β-sheet structure (dotted arrow). As the droplet matures, β-sheet conversion is initiated from IDRs, forming an insoluble gel. Finally, the entire molecule is converted to β-sheet-rich structure and oligomerized, resulting in proteinase-resistant gel. Yellow green: PEG fraction and PEG molecule. Light blue: Dextran fraction. Red: IDRs of rPrP. Dark blue: constructed region of rPrP including 3 α-helices. Purple dots: kosmotropic anions. Arrows: β-sheet structure.

Although our experimental conditions were highly artificial, it is still worthy to consider the possibility that the liquid-liquid phase separation of PrP^C^ could occur *in vivo*. Liquid-liquid phase separation is a phenomenon that was initially reported for intracellular proteins but later, proteins associated with the cell membranes were also shown to undergo liquid-liquid phase separation. Recently, it has been reported that zona occludens, a membrane-associated scaffolding protein, underwent liquid-liquid phase separation to form functional tight junctions between cells (38), suggesting that the protein complex attached to the membrane certainly has the properties of liquid. PrP^C^ is anchored to the cell membrane and can interact with various macromolecules, including proteins, RNA, and lipids (39-41). These macro-biomolecules are intertwined to each other and might drive liquid–liquid phase separation of the membrane protein. Interestingly, it has been demonstrated that Aβ-oligomers and DNA-aptamers drive the liquid-solid phase transition of rPrP through the interaction of amino acid residues around 90-120 (42,43). Furthermore, both full-length prion protein and prion protein peptide (amino acid, 23-144) could form proteinase resistant, spherical-ellipsoid aggregates that grow as amyloid fibrils by the addition of detergent or polysaccharides, and thus supporting our hypothesis that liquid-solid phase transition is associated with prion diseases (44-46). Thus, our results suggest that PrP can autonomously form a liquid phase triggered by the interaction between the N-terminal region under certain conditions. These phenomena possibly happen in the following situation: when PrP^C^ are packed in endocytic vesicles, for example exosome, or PrP^C^ and other proteins complexes are crowded at the cell membrane, such as lipid raft. In such situations, interactions between multivalent and flexible IDR of PrP might further condense the molecules, restraining their motions and directions. This might also potentially be a mechanism for the efficient propagation of PrP^Sc^ *in vivo* without any mechanical agitation. Taken together, microenvironments *in vivo* with high concentrations of kosmotropic anions may drive liquid-liquid phase separation of PrP^C^, leading to spontaneous intra- and/or extracellular PrP-amyloid formation. Further experiments using cell culture and *in vivo* imaging are needed to elucidate whether PrP^C^ can undergo liquid-liquid phase separation *in vivo*.

## Conclusions

Liquid-liquid phase separation of full-length rPrP using ATPS was demonstrated. The droplets of rPrP appeared only at the interface between PEG and dextran. The N-terminal region of prion protein (amino acids 23-89) and kosmotropic anions in neutral pH were essential for this reaction. Furthermore, the liquid-solid phase transition was found to be accompanied by β-sheet transition, resulting in proteinase K-resistance. These results suggest that spontaneous conversion of PrP^C^ to PrP^Sc^ amyloid may be promoted by liquid-liquid phase separation at the interface of biopolymers on the cell surface. Because of the limitations of our *in vitro* study, further approaches are required to elucidate whether the liquid-liquid phase separation of PrP^C^ on the cell surface can be provoked.

## Acknowledgments

We would like to thank Atsuko Matsuo, Hirono Nakata, and Ren Matsushima for technical assistance, and Toshiyuki Tsurumoto and Ryoichi Mori for polarized light microscopy observations. We would like to thank Editage (www.editage.com) for English language editing. This research was supported by JSPS KAKENHI Grant numbers JP19K22600 and Sasakawa Scientific Research Grant from The Japan Science Society.

## CRediT authorship contribution statement

Hiroya Tange: Conceptualization, Methodology, Validation, Formal analysis, Investigation,

Data Curation, Writing - Original Draft, Review & Editing, Visualization, Funding acquisition

Daisuke Ishibashi, Takehiro Nakagaki, Hiroki Ozawa: Resources, supervision

Yuji O. Kamatari: Data Curation and formal analysis

Yuzuru Taguchi: Writing-Review & Editing, Supervision

Noriyuki Nishida: Writing-Review & Editing, Supervision, Management, and Coordination

Responsibility for Research Activity Planning and Execution, Funding Acquisition

## Legend for supplemental Figures

**Sup. Fig. 1.**
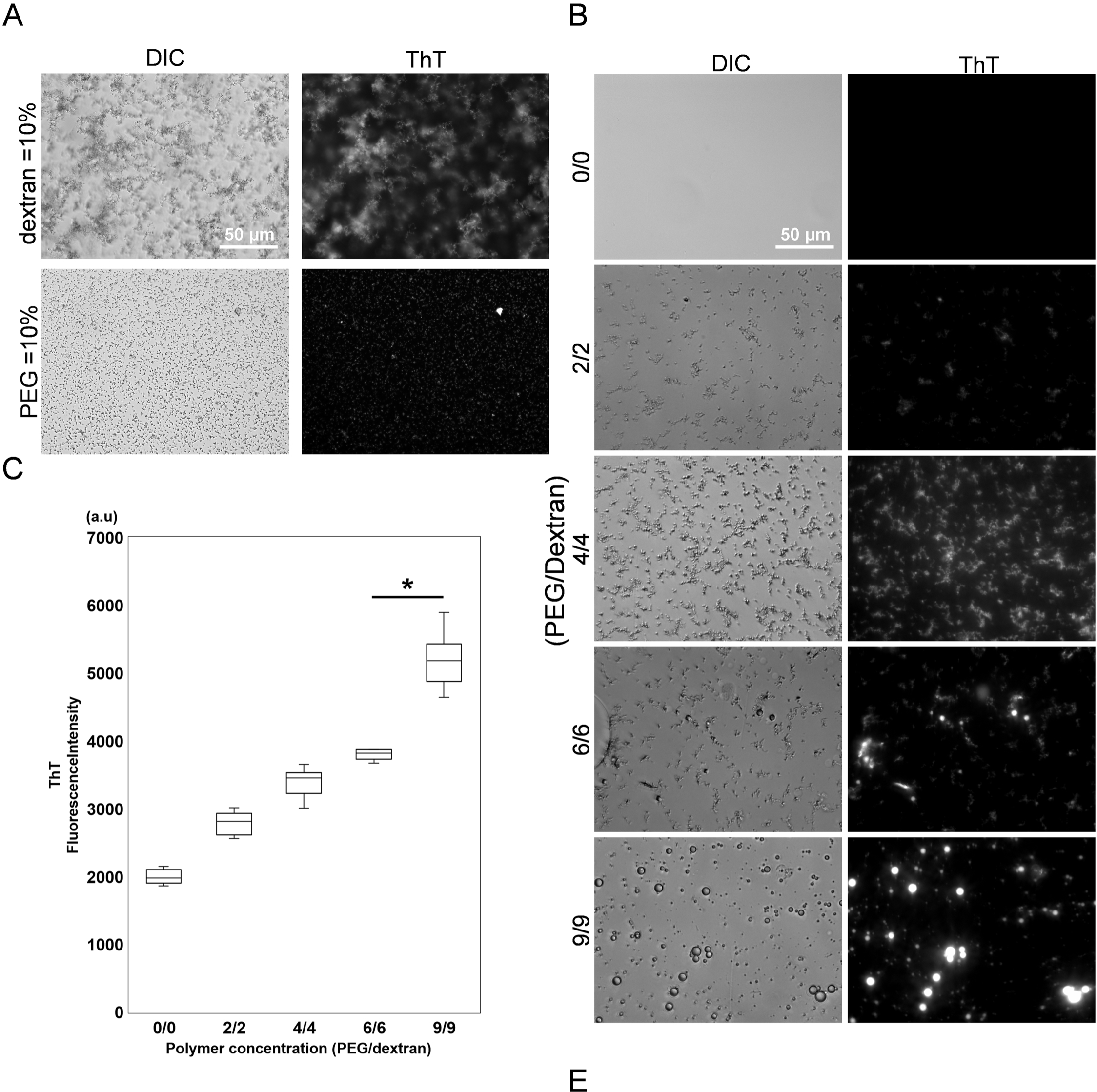
rPrP undergoes liquid phase separation above the binodal curve of ATPS. (A) DIC and fluorescence microscopic images of rPrP (10 µM) mixed with 10% dextran (Top) or 10% PEG (bottom) with 120 mM thiosulfate sodium. rPrP salted out as ThT-positive aggregates. (B) DIC and fluorescence microscopic images of the bottom of the well after 24 h of incubation are shown in Fig. 1B. ThT-positive amorphous aggregation precipitated in 2% –4%/2 – 4%, whereas a puddle-like droplet (white arrowhead) and amorphous aggregation coexisted with spherical-shaped precipitates in 6%/ 6%, and only spherical precipitates were observed in the 9%/ 9% condition. (B) Box-and-whisker plots for ThT fluorescence intensity of each polymer concentration after 24 h of incubation. The upper and lower whiskers represent the full range of values. The central horizontal lines indicate median values. Boxes illustrate the ranges between the lower and upper quartiles. N=9. *P<0.0001. Statistical analysis was performed with one-way ANOVA, followed by the Tukey-Kramer test.

**Sup. Fig. 2.**
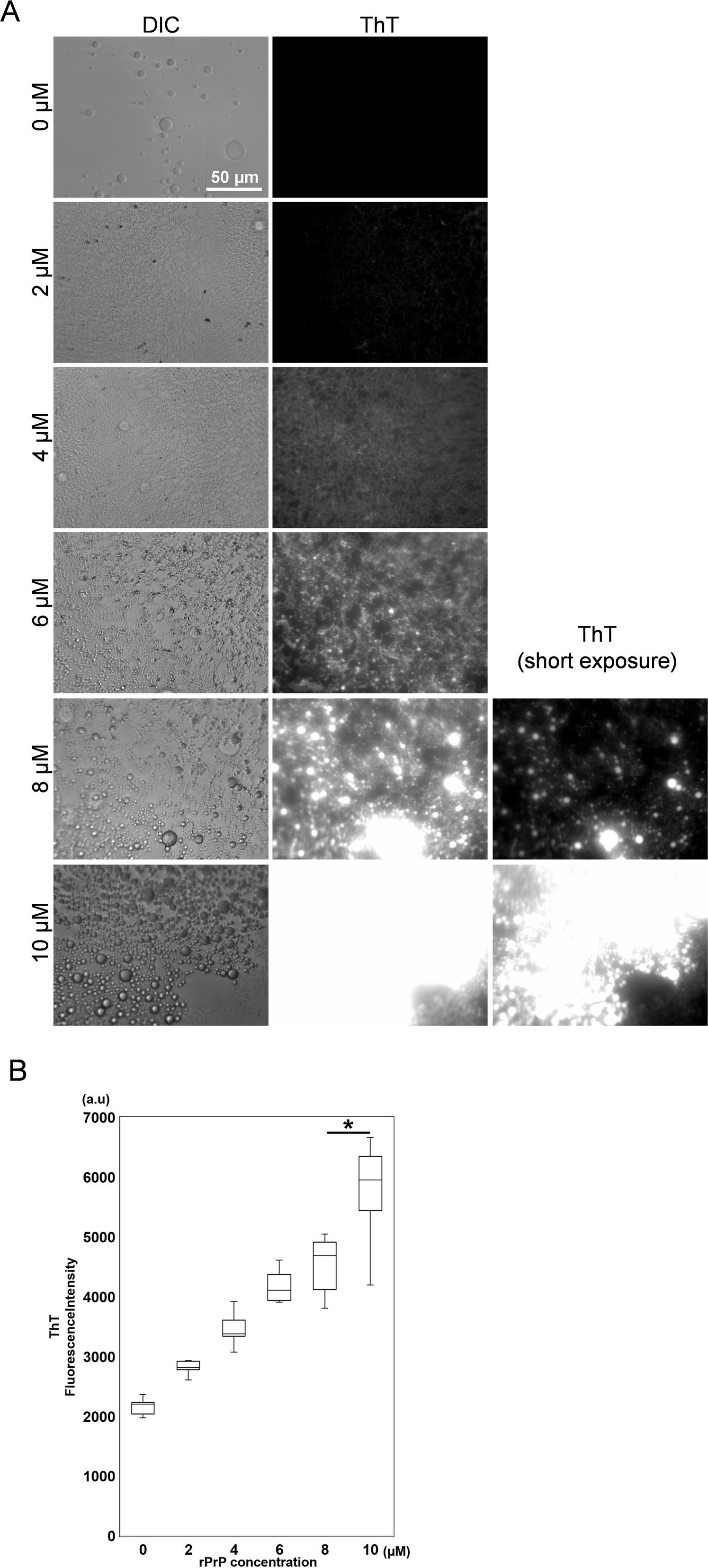
Droplet formation efficiency corresponding to rPrP concentration. (A) DIC and fluorescence microscopic images of rPrP at each concentration after 24 h of incubation. (B) Quantification of ThT fluorescence intensity for each rPrP concentration after 24 h of incubation. The upper and lower whiskers represent the full range of values. The central horizontal lines indicate median values. Boxes illustrate the ranges between the lower and upper quartiles. N=9. *P<0.0001. Statistical analysis was performed with one-way ANOVA, followed by the Tukey-Kramer test.

**Sup. Fig. 3.**
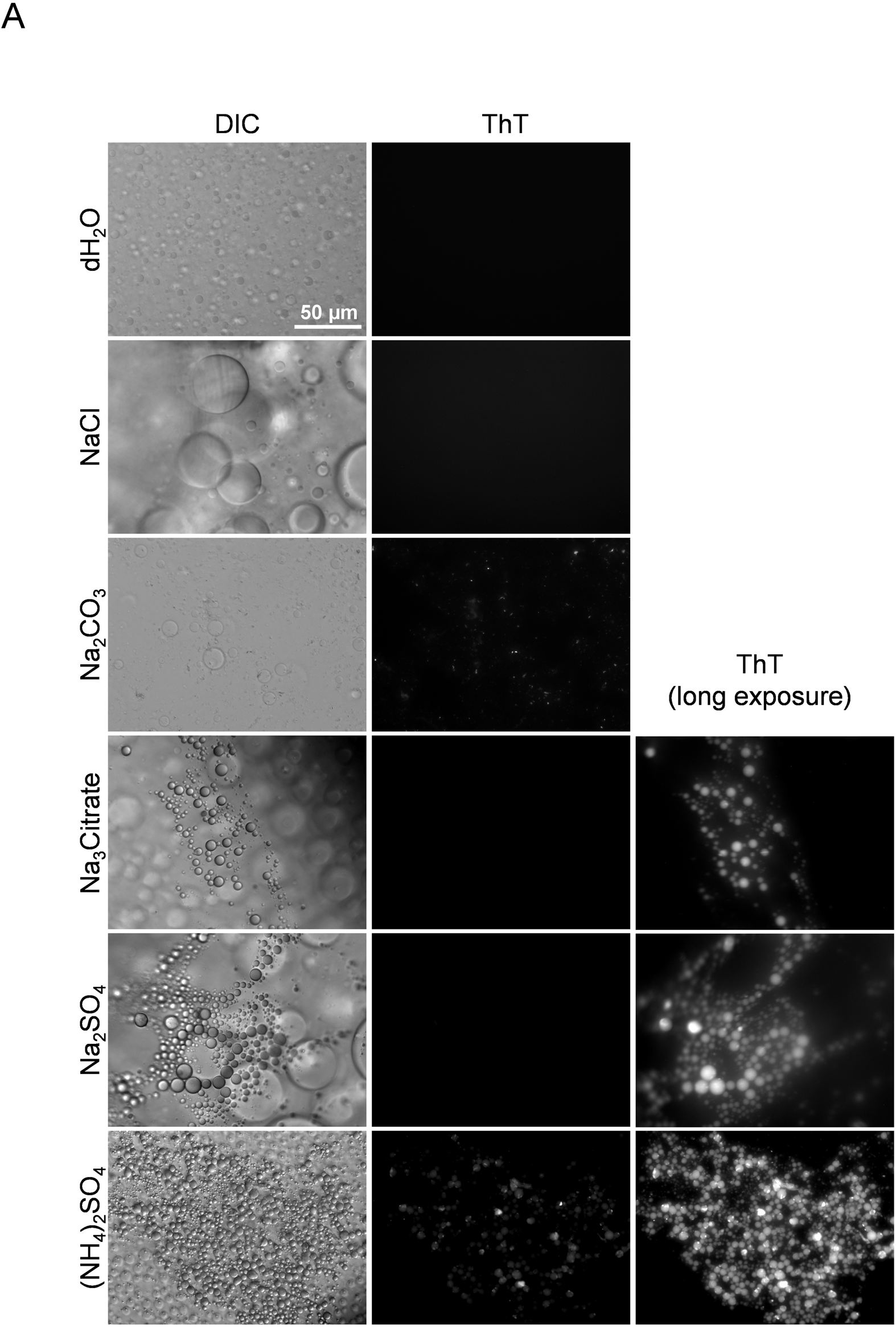
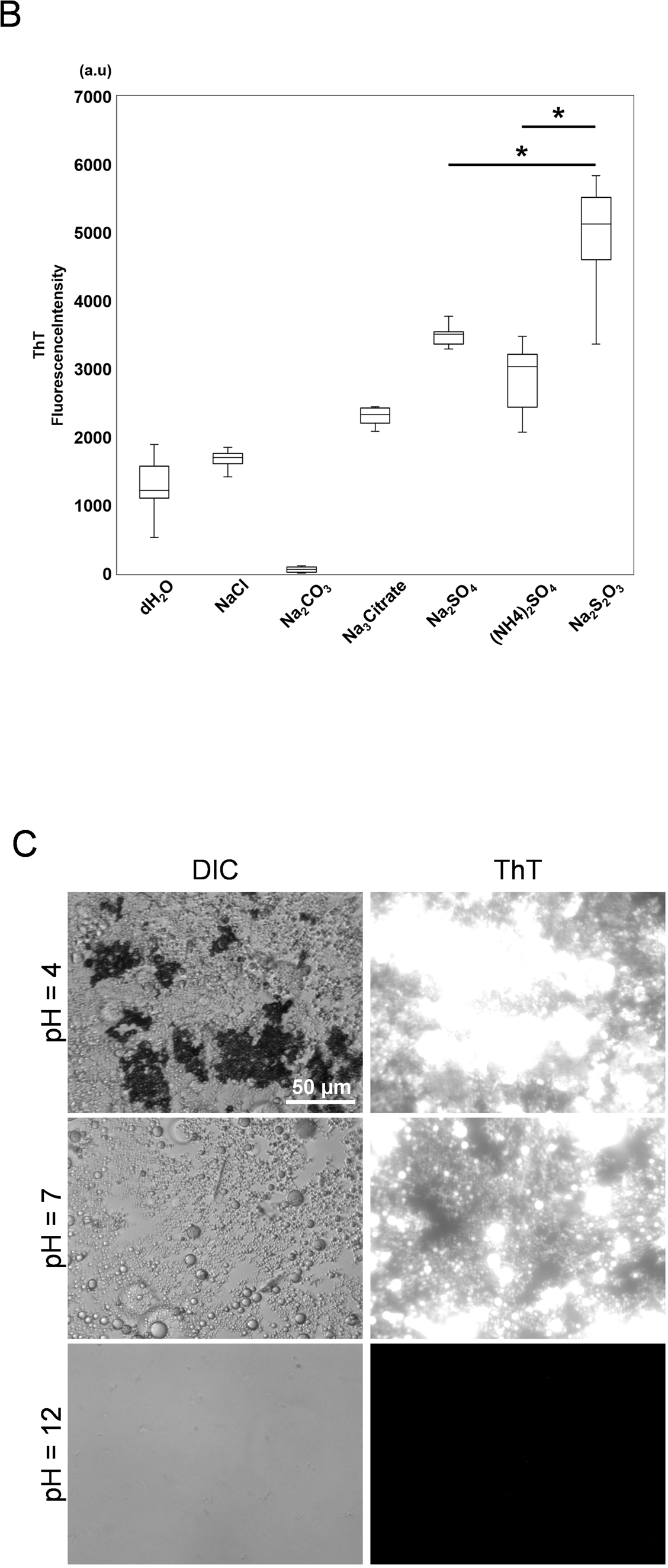
Effect of salts and different pH values on droplet formation. (A) DIC and fluorescence microscopic images of the interface of wells after 24 h of incubation with various salts. Left: DIC. Middle: fluorescence microscopic images Right: Acquired with high exposure. (B) Quantification of the fluorescence intensity according to the salts. The upper and lower whiskers represent the full range of values. The central horizontal lines indicate median values. Boxes illustrate the ranges between the lower and upper quartiles. N=9. *P<0.0001. Statistical analysis was performed with one-way ANOVA, followed by the Tukey-Kramer test. (C) DIC and fluorescence microscopic images at each pH after 24 h of incubation.

**Sup. Fig. 4.**
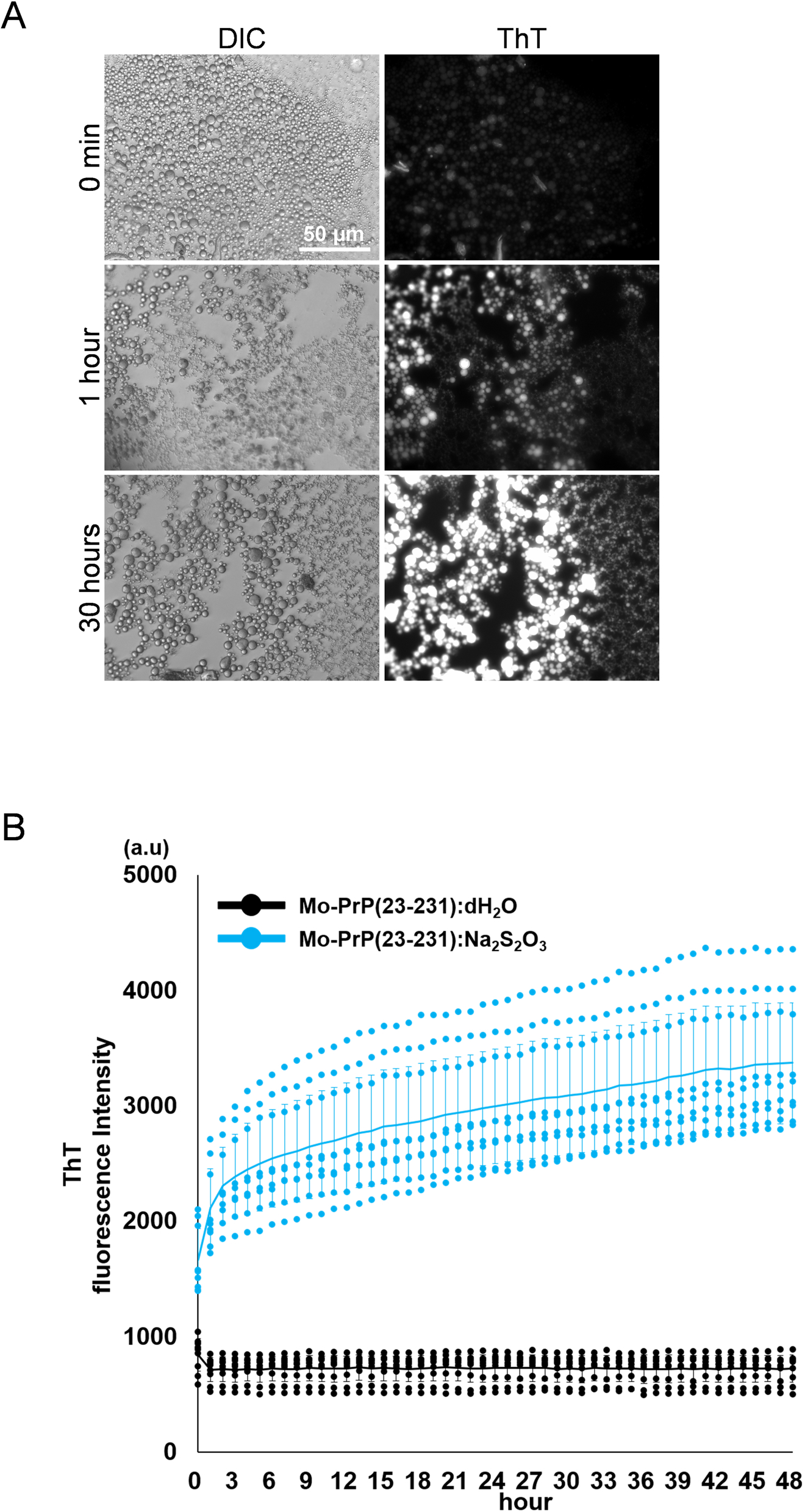
Mouse prion protein behaves similar to human prion protein. (A) DIC and fluorescence microscopic images of Mo-rPrP (23-231): dH2O and Na_2_S_2_O_3_ at 0 min, 1 h, and 30 h. (B) ThT fluorescence intensity of Mo-rPrP (23-231): dH_2_O and Mo-rPrP (23-231):Na_2_S_2_O_3_ measured after 48 h. Each dot represents a value measured in 3 independent experiments. Line indicates the average of each group. Error bars represent the standard deviation. N=9.

## Graphical abstract

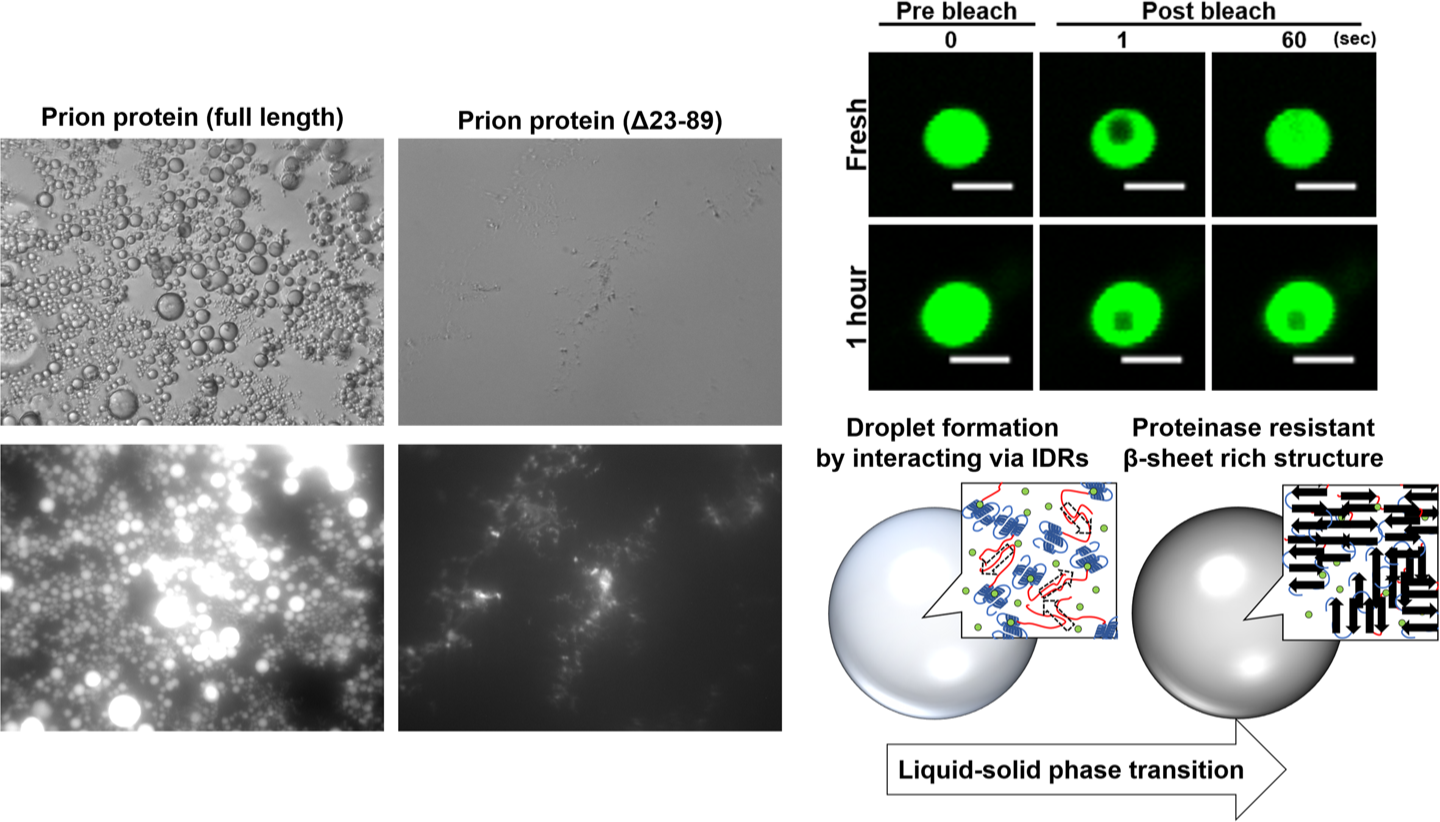

